# The distribution of fitness effects of spontaneous mutations in *Chlamydomonas reinhardtii* inferred using frequency changes under experimental evolution

**DOI:** 10.1101/2021.09.29.462298

**Authors:** Katharina B. Böndel, Toby Samuels, Rory J. Craig, Rob W. Ness, Nick Colegrave, Peter D. Keightley

## Abstract

The distribution of fitness effects (DFE) for new mutations is fundamental for many aspects of population and quantitative genetics. In this study, we have inferred the DFE in the single-celled alga *Chlamydomonas reinhardtii* by estimating changes in the frequencies of 254 spontaneous mutations under experimental evolution and equating the frequency changes of linked mutations with their selection coefficients. We generated seven populations of recombinant haplotypes by crossing seven independently derived mutation accumulation lines carrying an average of 36 mutations in the homozygous state to a mutation-free strain of the same genotype. We then allowed the populations to evolve under natural selection in the laboratory by serial transfer in liquid culture. We observed substantial and repeatable changes in the frequencies of many groups of linked mutations, and, surprisingly, as many mutations were observed to increase as decrease in frequency. We developed a Bayesian Monte Carlo Markov Chain method to infer the DFE. This computes the likelihood of the observed distribution of changes of frequency, and obtains the posterior distribution of the selective effects of individual mutations, while assuming a two-sided gamma distribution of effects. We infer that the DFE is a highly leptokurtic distribution, and that approximately equal proportions of mutations have positive and negative effects on fitness. This result is consistent with what we have observed in previous work on a different *C. reinhardtii* strain, and suggests that a high fraction of new spontaneously arisen mutations are advantageous in a simple laboratory environment.

## Introduction

Understanding the nature of genetic variation for fitness requires an understanding of the origin of that variation from mutation. The distribution of fitness effects of mutations (DFE) describes the frequencies of mutations with differing magnitudes of effects, and is fundamental for many topics in evolutionary genetics, including the maintenance of genetic variation, the nature of genetic variation for quantitative traits and the genetic basis of adaptive evolution. The DFE specifies the relative frequencies of advantageous and deleterious mutations and the contributions of mutations with small and large effect sizes to fitness change and genetic variation. The DFE appears, for example, in the nearly neutral model of molecular evolution (Ohta 1973), which posits that patterns of molecular variation and between species change can be explained by mutations that have fitness effects close to 1/*N*_*e*_ (where *N*_*e*_ is the effective population size); this is a very small fitness effect for species with typically large *N*_*e*_.

The DFE can be inferred experimentally or by statistical analysis of the frequencies of nucleotide variants at polymorphic sites (Eyre-Walker and Keightley, 2007). In the latter approach, the DFE for deleterious amino acid-changing mutations can be estimated by analysis of the site frequency spectrum (SFS) for nonsynonymous variants under the assumption that their distribution of effects follows a pre-specified distribution, such as a gamma distribution. When the SFS is combined with divergence data from another species the frequency and effects of advantageous nonsynonymous mutations can also be estimated (Eyre-Walker and Keightley 2009; Tataru et al. 2017). Analysis of the SFS has been applied to genomic data from a wide range of taxonomic groups. Estimated DFEs for deleterious mutations are invariably strongly leptokurtic (L-shaped), the shape of the distribution varying between taxonomic groups (Chen et al 2017), and there is usually a strong nearly neutral component. Analysis of the unfolded SFS suggests that at most a few percent of mutations are advantageous (Keightley et al 2016). Inferring the DFE using standing variation within a population is relevant to the fitness effects of mutations in nature, but does not capture strongly positively or strongly negatively selected mutations, because these tend not contribute to standing nucleotide variation.

Inference of the DFE via experimental manipulation can be applied to mutations engineered in specific genes or at random genomic locations. One approach, which has similarities to the one described here, estimates the selection coefficients of induced mutations by tracking their frequency change over time under experimental evolution. This was pioneered by McDonald et al (2016), who used deep sequencing to measure frequency changes of newly arisen mutations under experimental evolution in *Saccharomyces cerevisiae*. These frequency changes were used to estimate mutational effect sizes for fitness under adaptation, although McDonald et al (2016) did not infer the full DFE. More recently, Flynn et al (2020) used “deep mutational scanning” of yeast strains to infer the DFE for mutations in the Hsp90 gene. By competition between strains followed by deep sequencing, Flynn et al quantified growth effects of many single codon changes encoding amino acid variants under standard environmental conditions and under five stress conditions. In standard conditions, the DFE is a leptokurtic distribution, containing a small proportion of beneficial mutations. However, the proportion of beneficial mutations was substantially higher in non-standard or stressful environments, especially high temperature and salt.

The DFE can also be inferred for mutations induced at random locations in the genome. For example, Johnson et al (2019) estimated the DFE for transposable element mediated insertion mutations in yeast by measuring mutation frequency changes under adaptation over time. Notably, lines with the highest initial fitness appeared to suffer the greatest fitness consequences from *de novo* insertion mutations.

Previously, we have inferred the DFE for spontaneous mutations in the single-celled green alga *Chlamydomonas reinhardtii* using growth rate as a fitness measure (Böndel et al 2019). We crossed mutation accumulation (MA) lines of the CC-2931 strain that had randomly accumulated spontaneous mutations for many generations with a mutation-free ancestral strain. We then measured growth rate and determined the genotypes of many recombinant lines carrying random combinations of mutations. We developed a Bayesian MCMC approach to estimate parameters of a DFE, which also enabled us to extract the estimated effects of individual mutations. This suggested a highly leptokurtic DFE, with a surprisingly high proportion of mutations (about 50%) increasing growth rate.

Here, we cross MA lines of *C. reinhardtii* of a different strain (CC-2344) to that studied by Böndel et al (2019), which had undergone approximately 1,000 generations of spontaneous mutation accumulation, with a mutation-free ancestral strain in order to generate many recombinants with different combinations of mutations. Rather than assaying individual recombinants, as in our previous experiment, we allow pools of recombinant haplotypes to compete against one another in an experimental evolution setting and measure changes of mutation frequency by deep sequencing. We develop a new MCMC approach to infer the DFE based on changes in frequency of linked mutations. Using this new experimental approach and a different strain, we infer that a surprisingly high proportion of mutations increase fitness in the standard laboratory environment.

## Materials and Methods

### Mutation accumulation lines and compatible ancestor

We studied seven *C. reinhardtii* MA lines (L06, L09, L10, L12, L13, L14, L15) derived from strain CC-2344 (isolated in Pennsylvania, USA, in 1988) produced by Morgan et al. (2014) and sequenced as described by Ness et al. (2015). The MA lines and their ancestral strain are of the same mating type (mt+) and will not mate with one another, so we first produced a “compatible ancestor” to which the MA lines could be crossed. This was done by backcrossing CC-2344 to a mt-strain (CC-1691) for 16 generations with the aim of producing a strain nearly identical to CC-2344, with the exception of the region surrounding the mating type locus on chromosome 6. Genome sequencing of the compatible ancestor (using the method of Ness et al. 2015) showed that this was accomplished successfully (Figure S1).

### Generation of populations of recombinants

To produce the starting populations of the experimental evolution experiments (designated *t*_0_), we generated recombinant populations by mating each MA line with the compatible ancestor. First, we grew each MA line in Bold’s medium (Bold 1942) under standard conditions (23°C, 60% humidity, constant white light illumination) while shaking at 180 rpm to obtain a culture of 30 ml. Three lines had poor growth (L09, L12, and L15), so this procedure was done twice in order to obtain sufficient cell material. After three days, cultures were transferred to 50 ml falcon tubes, centrifuged at 3250 g for 5 minutes and the supernatant removed. Nitrogen-free conditions are required to trigger mating in *C. reinhardtii* (Sager and Granick 1954), so we washed each cell pellet with 30 ml nitrogen-free Bold’s medium, mixed, centrifuged at 3250 g for 5 minutes and removed the supernatant. We then resuspended the washed cell pellet in 45 ml nitrogen-free Bold’s medium. The same procedure was done for the compatible ancestor.

To carry out the matings, we mixed 15 ml of the resuspended MA line cell culture with 15 ml of the resuspended compatible ancestor cell culture in a 50 ml falcon tube. The mixtures were then incubated under standard growth conditions for 7 days in a slanting position (in order to increase the surface area) until zygote mats had formed at the surface. The remaining 30 ml of each culture served as control to allow the detection of mating failures (see below). Zygote mats were then transferred to a fresh 50 ml falcon tube containing 30 ml nitrogen-free Bold’s medium and incubated in the dark under standard growth conditions for 5 days to allow the zygotes to mature. To kill any vegetative cells still associated with the zygote mats, we froze the zygote cultures and kept them at −20 °C for 5 hours. After thawing at room temperature for approximately 1 hour, we transferred the zygote cultures to 500 ml conical flasks and added 30 ml of Bold’s medium containing twice the standard concentration of nitrogen and 60 ml of standard Bold’s medium to obtain a total volume of 120 ml with standard nitrogen concentration. The flask was then incubated under standard growth conditions while shaking at 250 rpm until zygotes had germinated (Figure 1A). This germination cell culture was then used to start the experimental evolution experiment. The control cultures were incubated as described for the zygote cultures and if any growth was visible after the freezing, the respective zygote culture would have been discarded and the whole procedure repeated, but no such instances were observed.

**Figure 1.**
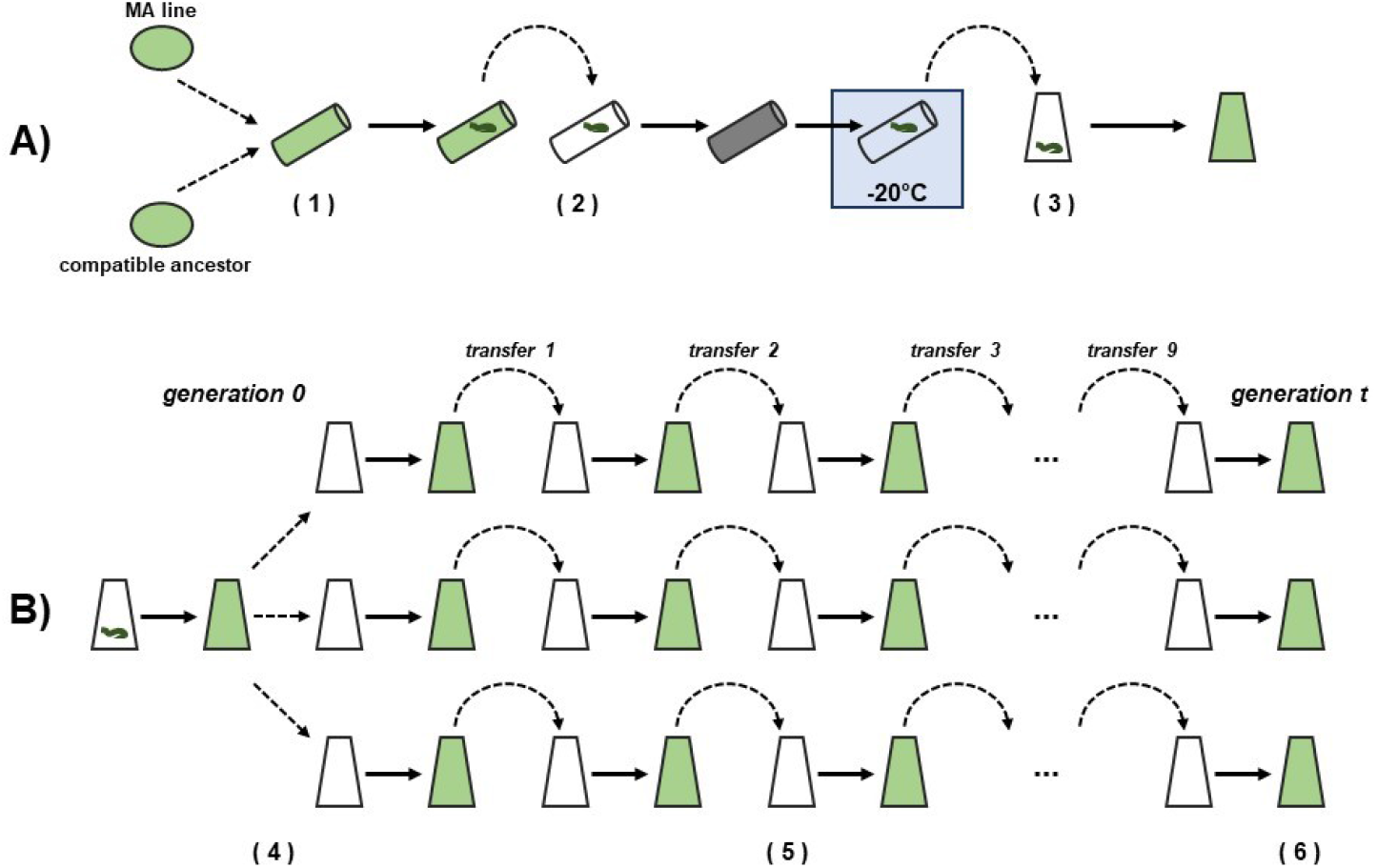
Schematic overview of the experimental procedure. A) Matings. Cell cultures of the MA line and the compatible ancestor were mixed in a falcon tube and incubated until zygote mats had formed (1). The zygote mats were transferred to new falcon tubes, incubated in the dark to allow zygote maturation and then frozen at −20°C to kill off any vegatative cells (2). The culture with the matured zygote mat was transferred to a conical flask and incubated until zygotes germinated (3). B) Serial transfers. The germination culture was grown up until it was dense enough to start the three replicates and have enough cell material for DNA extraction. The three replicates were started with 1.2 ml of the original germination culture (4). Every three to four days 1.2 ml were transferred to fresh conical flasks and 2 ml of the remaining culture was used to measure OD (5). After nine transfers the end of the experiment was reached and the cells were collected for DNA extraction (6).

### Experimental evolution

We grew three replicate cell cultures from each MA line x compatible ancestor cross until approximately 60 generations had been reached (Figure 1B). We designate this time point *t*_*t*_. For L06 and L09 this took 42 days and for L10, L12, L13, L14, and L15 this took 46 days. Each replicate was grown in a volume of 120 ml in a 250 ml conical flask under standard growth conditions while shaking at 250 rpm. We started each replicate with 1.2 ml of the germination cell culture and 118.8 ml Bold’s medium. By using standard Bold’s medium we ensured that the recombinants could not mate and grew entirely vegetatively. The remaining cells from the germination cell cultures were collected and frozen at −70°C for sequencing. We then transferred 1.2 ml of each culture to a fresh flask containing 118.8 ml Bold’s medium on a three or four day cycle, or in the case of transfers 3, 6 and 9, over seven days in order to maximise biomass for DNA extraction and sequencing. In order to determine the number of serial transfers at which approximately 60 generations had been reached, optical density (OD) at 600 nm of the cultures was measured at the end of each transfer growth period. We then calculated the generation time as follows: *t = (*log *N*_*t*_ *–* log *N*_*0*_*) /* log 2, where *N*_*t*_ is the measured OD of the culture at the end of the growth period and *N*_*0*_ the calculated OD after dilution of the previous transfer at the beginning of the growth period. After nine transfers (approximately 60 generations) we collected cells for sequencing and froze the pellets as described for the time point 0 samples.

### Sequencing and sequence data processing

Genomic DNA was obtained from time point 0 (*t*_*0*_) and from each of the three *t*_*t*_ replicates for each MA line recombinant population by phenol-chloroform extraction (Ness et al 2012). DNA samples were sequenced on an Illumina Hiseq4000 platform by BGI Hong Kong with 150 bp paired-end reads to an average sequencing depth of 520.7x and 221.6x for time 0 and *t*, respectively. Fastq reads were mapped to the *C. reinhardtii* reference genome (strain CC-503; version 5; Merchant et al. 2007) with bwa-mem (Li and Durbin 2009) and duplicate reads were removed with MarkDuplicates using picard tools. The four bam files of each MA line (time 0, and the three replicates of time *t*) were merged with samtools (Li et al. 2009) into a single bam file. The 1,000-bp regions surrounding the locations of the mutations of interest (500 bp before and 500 bp after each mutation) were then realigned with HaplotypeCaller of GATK (McKenna et al. 2010, DePristo et al. 2011) to allow more accurate mapping of the mutations and more accurate allele frequency estimation. The realigned bam files were then split into the individual samples with SplitSamFile of GATK (McKenna et al. 2010, DePristo et al. 2011). Samtools was then used to create pileup files from the realigned bam files using the mpileup command. Allele frequencies for the mutations of interest were then calculated with custom Perl scripts.

Mutations were classified as noncoding, synonymous or nonsynonymous relative to the v5.3 reference genome annotation. To assess the functional effects of mutations, SnpEff (Cingolani et al. 2012) was run using the pre-build *C. reinhardtii* annotation and with default parameters.

It has recently been suggested that the v5 reference genome contains some misassemblies (Salomé & Merchant, 2019; Craig et al. 2021). Since misassemblies could introduce incorrect linkage relationships between mutations, we lifted over mutation coordinates to the highly contiguous Nanopore-based assembly of the strain CC-1690 (O’Donnell et al., 2020), which is identical-by-descent >95% of its genome with the original reference strain (Gallaher et al. 2015). The v5 and CC-1690 assemblies were aligned with Cactus (Armstrong et al. 2020) with divergence between the genomes arbitrarily set at 0.004 and otherwise default parameters. Liftover was then achieved from the resulting alignment using halLiftover (Hickey et al. 2013). Lifted over coordinates were used for the MCMC analysis described below.

### Repeatability of mutation frequency between replicates

We estimated the repeatability of mutation frequency among the three replicate populations by partitioning the total variation in allele frequencies into three components: the variance in frequency among different MA line x compatible ancestor crosses (*V*_*MA*_), the variance in frequency among mutations (*V*_*M*_) and the residual variation due to differences among the three replicate measures for each mutation (*V*_*E*_). Variance components were estimated using the Lmer function in R, and repeatability was calculated as *V*_*M*_/(*V*_*M*_ + *V*_*E*_).

### *C. reinhardtii* genetic map

A genetic map is required in the MCMC analysis described below, which calculates frequency changes of linked mutations. We assume an overall genome-wide average rate of recombination, obtained from two published crosses between *C. reinhardtii* strains CC2935 x CC2936 and CC408 x CC2936, which together provide an estimate of 1cM per 87,000 bases (Liu et al 2018). A third cross reported by Liu et al. (2018) (CC124 x CC1010) that has a *c*.10-fold lower marker density was not included in our calculations.

Our analysis required the location of the individual mutations on a genetic map. Lui et al. (2018)’s study does not provide sufficient resolution for this, but higher resolution estimates of the rate of recombination are available from a study of linkage disequilibrium in natural populations (Hasan and Ness 2020). We used these estimates to adjust for variation in the rate of recombination among chromosomes, i.e. assumed a uniform rate of recombination per chromosome. Longer chromosomes have lower recombination rates, presumably due to a requirement for a minimum of 1 chiasmata per chromosome per meiosis (Figure S2).

We used estimates of the population scaled recombination rate from Hasan and Ness (2020) to adjust the rate of recombination estimated in the Liu et al. (2018) crossing experiment in order that the inverse rate of recombination (*y*_*i*_, bp/cM) increases linearly by 0.000616 per 1 base pair increase in chromosome length (*x*_*i*_) (Figure S2), while keeping the overall average recombination unchanged, i.e.,

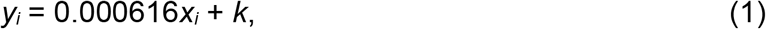

where

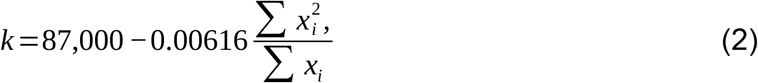

(see supplementary text A).

### Computation of expected allele frequencies after experimental evolution

In this section we describe the computation of the expected frequencies of the mutations after one generation of recombination followed by *t* generations of experimental evolution, which are used in likelihood calculations and Bayesian inference. We assume that allele frequencies of each mutation are 0.5 after crosses between MA lines and their ancestor, and that changes of allele frequency then occur deterministically and independently among chromosomes, but that genetic linkage of mutations on the same chromosome leads to non-independent allele frequency changes under selection. Based on the assumed genetic map (see above), we first computed the expected frequencies of the *n* possible haplotypes generated by one round of meiosis involving an ancestral chromosome and a chromosome from a MA line in the absence of selection. In the model of experimental evolution following the cross, mutations have selection coefficients, *s*_*j*_, from which the overall fitness (*w*_*i*_) of haplotype *i* carrying *m*_*i*_ mutations can be computed under multiplicative selection:

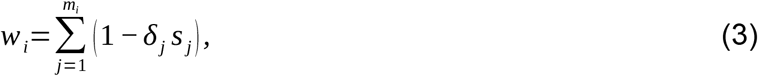

where *δ*_*j*_ takes the value 1 or 0 if the haplotype carries the mutant or wild type allele, respectively, for mutation *j*. The selection coefficients (*s*) are parameters of the model whose values are changed during MCMC runs. These and other such parameters are listed in Table 1.

**Table 1.** Parameters of the model.

| Parameter | Definition |
| --- | --- |
| $s_j$ | Selective effect of mutation $j$. |
| $\delta_{jk}$ | Deviation of the mean frequency of mutation $j$ replicate $k$ about its expectation. |
| $\alpha$ | Vector of scale parameters of the distribution of effects of mutations, elements 0 and 1 are for positive and negative effects, respectively. |
| $\beta$ | Vector of shape parameters of the distribution of effects of mutations. |
| $q$ | Frequency of positive effect mutations. |

Let the frequency of haplotype *i* at generation *t* = *π*_*i,t*_. We then calculated the expected frequency of each haplotype after *t* generations of natural selection by iterating equation (4) for *t* generations:

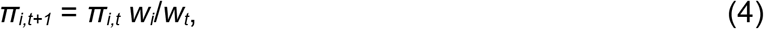

where *w*_*t*_ is the mean fitness of the *n* haplotypes at generation *t*. It is then straightforward to compute the expected frequencies of the individual mutations (*p*) at generation *t*.

### Computation of likelihood

The log likelihood was the sum of three terms, the first involving the mutation frequencies after selection (*p*), the second involving mutation effects (*s*), which were assumed to be gamma distributed, and the third involving the total number of negative effect mutations (*n*_0_) *versus* the total number of positive effect mutations (*n*_1_), which were assumed to be sampled from a binomial distribution with mean *q* (the frequency of positive effect mutations, a parameter of the model, Table 1). Likelihood was computed assuming independence among the *c* chromosomes:

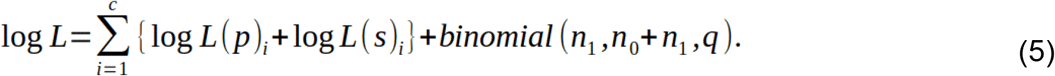

The log likelihood for the term involving the mutation frequencies on a chromosome (log *L*(*p*)_*i*_) was computed assuming that allele frequencies of each of the *r* experimental evolution replicates are sampled from a log-normal distribution with mean *p*_*j*_ and standard deviation *σ*_*δ*_, and that numbers of mutant and wild type reads for a replicate are sampled from a binomial distribution with mean *p*_*j*_ + *δ*_*jk*_:

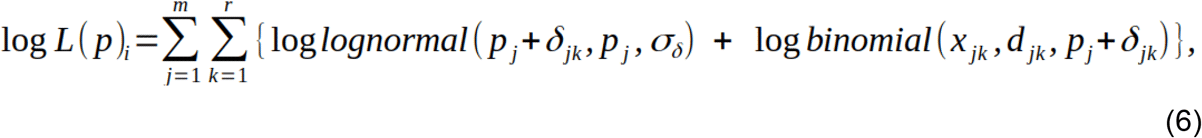

where *δ*_*jk*_ are variables in the model that allow the allele frequency for each replicate (*k*) to be different from the overall expected frequency under selection for that mutation, and *x*_*jk*_ and *d*_*jk*_ are the numbers of mutant reads and the sequencing depth, respectively, for mutation *j* replicate *k*.

The log likelihood for the term involving the distribution of fitness effects of mutations (log *L*(*s*)_*i*_) on a chromosome was computed assuming that the fitness effects are drawn from gamma distributions, which can have different parameters for positive and negative effect mutations:

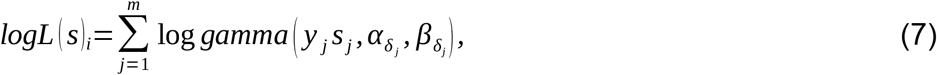

where gamma() is the gamma probability density function (PDF), *y*_*j*_ takes the value −1 or 1 if mutation *j*’s fitness effect is negative or positive, respectively, *δ*_*j*_ is an indexing variable that takes the value 0 or 1 if mutation *j*’s fitness effect is negative or positive, respectively, and 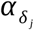and 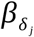 are elements of vectors **α** and **β**, both of dimension two, containing the scale and shape parameters, respectively, of the gamma distributions of fitness effects. Elements 0 and 1 of **α** and **β** contain parameters for negative and positive effect mutations, respectively.

### Priors

These were designed to be informative. The prior for fitness effects of mutations was a uniform distribution bounded by −1 and +1. The prior for the frequency of positive-effect mutations (*q*) was uniform in the range 0 and 1. Priors for the shape and scale parameters of the gamma distribution of effects of mutations and the parameters related to the log normal distribution (*σ*_*δ*_ and *δ*_*jk*_) were uniform in the range 0 to very large values.

### MCMC implementation

We used the Metropolis Hastings algorithm to sample from the posterior distributions of the parameters (Table 1), based on the product of the log likelihood of the data and priors (which were designed to be uninformative, see above). Briefly, there was a burn-in of 10^9^ iterations followed by sampling every 10^5^ iterations up to iteration 10^10^. Proposal deviates were sampled from normal distributions and added to the current parameter values. During the burn-in, the variance of a proposal distribution was either increased or decreased by a factor of 1.2 each iteration so that the average proportion of accepted proposals for each parameter was about 0.234. The mode of the posterior distribution was used as the parameter estimate and 95% credible intervals were computed based on ranked posterior values.

## Results

### Distribution of initial mutation frequency before experimental evolution

We crossed seven *C. reinhardtii* MA lines that had been independently derived from strain CC-2344 to a compatible ancestor of the same genetic background in order to generate populations of recombinants. In the generation following the cross, mutations are therefore expected to be at a frequency of 0.5. Recombinant haplotypes were then allowed to compete with one another in the absence of further recombination in standard laboratory conditions over the course of nine serial transfers. We sequenced samples from each MA line cross at the start (designated time 0) and end (designated time *t*) of experimental evolution in order to quantify changes in mutation frequency.

We identified 254 mutations in the seven MA lines, comprising 232 SNPs, 13 insertions and nine deletions (Table 2). Recombinant populations were sequenced at a high enough depth at *t*_0_ and *t*_*t*_ to allow accurate frequency estimation (Table 2). Sequencing depth varied among the mutations, but we did not observe a significant correlation between sequencing depth and mutation frequency (Figure S3; time *t*_0_: *r* = 0.0741, P = 0.236 and *t*_*t*_: *r* = 0.0244, P = 0.498; *r* = Spearman’s correlation). Therefore, we can discount any effect of sequencing depth on mutation frequency.

**Table 2.** Numbers of mutations in each MA line and sequencing depth statistics at the start and end of experimental evolution for the corresponding recombinant populations. Average sequencing depth across all mutations and replicates is shown along with the standard deviation and range in parenthesis.

| Line | Mutation type |  |  |  | Sequencing depth |  |
| --- | --- | --- | --- | --- | --- | --- |
| | SNP | INS | DEL | total | Time 0 | Time $t$ |
| L06 | 21 | 2 | 2 | 25 | 628.0 (262.5; 47, 1366) | 254.9 (117.2; 19, 665) |
| L09 | 13 | 0 | 2 | 15 | 353.6 (154.5; 100, 536) | 166.6 (84.1; 30, 306) |
| L10 | 45 | 2 | 0 | 47 | 513.2 (226.8; 41, 1252) | 216.4 (101.6; 10, 608) |
| L12 | 50 | 2 | 0 | 52 | 509.6 (253.0; 8, 1767) | 204.9 (134.6; 5, 1016) |
| L13 | 35 | 4 | 2 | 41 | 487.1 (170.4; 120, 1011) | 227.4 (85.6; 40, 526) |
| L14 | 33 | 1 | 2 | 36 | 580.3 (214.1; 94, 1250) | 263.6 (130.4; 23, 723) |
| L15 | 35 | 2 | 1 | 38 | 520.3 (177.1; 92, 1134) | 204.5 (71.9; 21, 437) |
| all | 232 | 13 | 9 | 254 | 520.7 (221.7; 8, 1767) | 221.6 (111.0; 5, 1016) |

The average mutation frequency at time 0 (*p*_0_) was 0.481, which is close to the expected value of 0.5. Initial frequency of the different mutations showed considerable scatter, however, since the standard deviation was 0.141, and there were also noticeable differences in the distribution of frequencies between the recombinant populations. Lines L12 and L13, for example, had a relatively broad range of initial frequencies centering around 0.5, whereas L9 and L10 had a narrower distribution, with a mean close to 0.5 and a few mutations with very low frequencies (Figure 2). Unexpectedly, in the majority of lines there were mutations at frequencies close to zero at time 0, and additionally the frequency of one mutation L15 was close to 1 at time 0 (Supplementary Figure S4). These extreme frequencies could be explained by natural selection changing mutation frequency in the generations of growth prior to the sequencing of the populations at time 0. This is corroborated by the presence of groups of linked mutations having similar frequencies at time 0 (Supplementary Figure S4).

**Figure 2.**
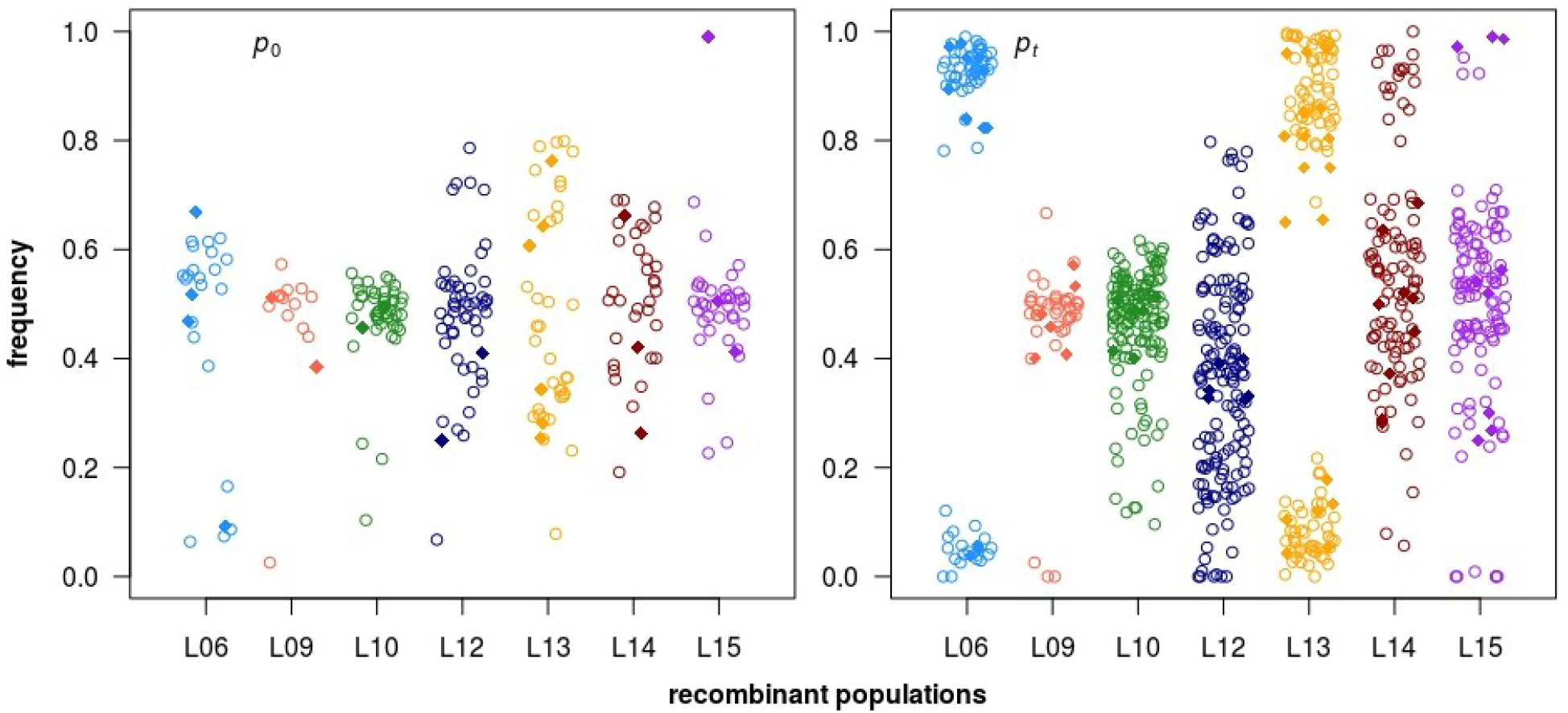
Mutation frequencies at the start (*p*_0_) and the end (*p*_*t*_) of experimental evolution of the seven recombinant populations. Mutational types are indicated with symbols: open circles - SNPs, closed diamonds - Indels.

### Mutation frequencies after experimental evolution

There were substantial changes in the frequencies of many mutations after experimental evolution (*p*_*t*_), but the magnitude and direction varied substantially among mutations (Figure 2). Mutation frequency was highly repeatable between the three replicates (*r* = 0.96), indicating that selection was the causal agent of frequency change.

The correspondence between mutation frequencies at the start and end of experimental evolution is shown in Figure 3. As previously mentioned, *t*_*0*_ frequencies are centred around 0.5 (see also Figure 2 and Figure S4), but in some cases *t*_*0*_ frequencies were substantially different from 0.5. In the majority of these cases, frequency usually became even more extreme in the same direction by time *t*. This suggests that natural selection changed mutation frequency in the generations after the cross before sequencing at *t*_*0*_, and selection also operated in the same direction under subsequent experimental evolution. There are exceptions, however, the most extreme of which correspond to points in the top-left and bottom-right quadrants of Figure 3. These are cases where the allele frequency first moved down (top-left quadrant) and then moved up or *vice versa*. A possible explanation for this behaviour is a change in the direction of selection, which would imply that the environmental conditions before and after sampling for time 0 had changed.

**Figure 3.**
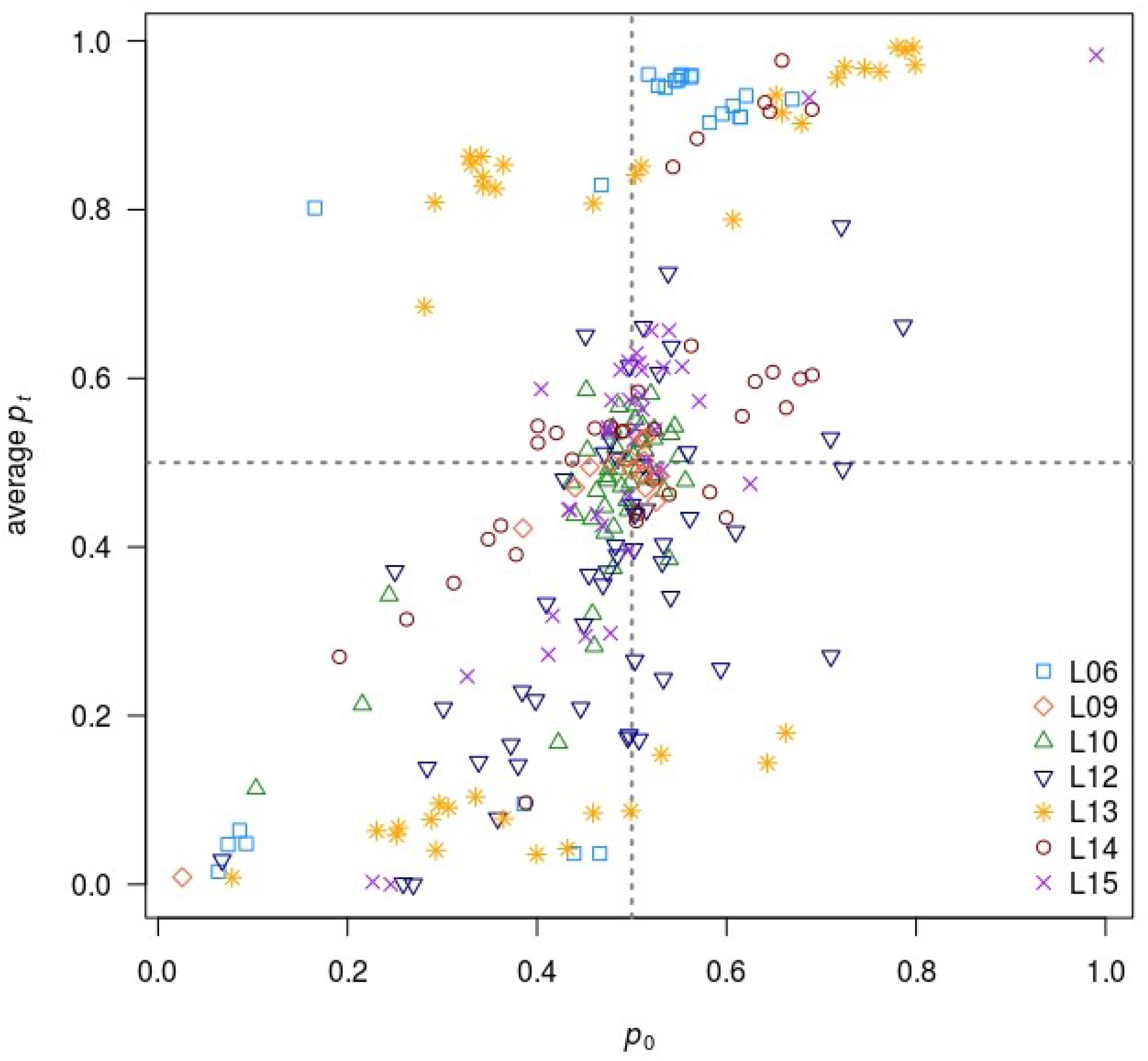
Mutation frequencies at the start of experimental evolution (*p*_0_) *versus* average mutation frequencies at the end of experimental evolution (*p*_*t*_). The different recombinant populations are indicated with different colours and symbols. The dotted lines represent the frequencies of 0.5.

Notwithstanding the unexpected cases mentioned above, there are other patterns apparent in the raw allele frequencies (Figure S4). To account for allele frequency variation among replicates (which are mostly very consistent), in the subsequent analysis, we modelled variance among replicates of the same cross by assuming a lognormal model for the environmental variance between replicates of the same cross. Linked mutations usually moved in frequency in the same direction.

### Estimation of the generation time

The number of generations over the course of experimental evolution were estimated based on OD measurements taken at each of the nine transfers. The mean number of generations across all recombinant populations was 59.8 with a standard deviation of 1.03. The estimated generation times were highly consistent between replicates (Figure S5, Table S1) and varied only slightly between recombinant populations from the different MA line x compatible ancestor crosses (Table S1) where L06 had the lowest number of generations (58.1) and L15 the highest (60.7).

### MCMC analysis to estimate the DFE

We proceeded to carry out the Bayesian MCMC analysis to estimate parameters of the DFE described in Methods, assuming that the number of generations of natural selection under experimental evolution for each MA line recombinant population *t* = 60, fitting three different two-sided gamma distributions of fitness effects: a distribution with the same shape and scale parameters for positive- and negative-effect mutations, a distribution with different scale parameters and the same shape parameters for positive- and negative-effect mutations, and a distribution with different scale and shape parameters for positive- and negative-effect mutations. In each case, after a burn-in period, the sampler appeared to have converged (Figure S6), and samples were drawn from the chain in order to obtain estimates of the posterior distributions of the various parameters.

### Model comparison

We used the convention that if BIC(model A)–BIC(model B) < −10, there is strong evidence in favour of model A over model B (Raffery 1995). The results (Table 2) therefore suggest that more complex models are not strongly favoured over the simplest two-sided gamma model, which has the same scale and shape parameters for positive- and negative-effect mutations.

**Table 2.** Models, their numbers of parameters related to the DFE and BIC values.

| ***Model*** | ***Number of parameters*** | ***BIC*** |
| --- | --- | --- |
| Two-sided gamma | 2 | 1.6 |
| Two-sided gamma, different means | 3 | 7.2 |
| Two-sided gamma, different means and shapes | 4 | 12.3 |

### Parameter estimates

We obtained estimates of parameter values based on the modes of the posterior distributions sampled from the MCMC chains. For the three different models evaluated, the parameter estimates are highly consistent (Table 3). The estimate of the proportion of positive-effect mutations (*q*) is close to 0.5 for each of the three models, and credible intervals are relatively narrow. This result is consistent with the observed bidirectional changes of mutation frequency. For each of the three models, estimates of the shape parameter of the distribution of effects are close to 0.5. This implies a highly leptokurtic distribution of fitness effect, in which the majority of effects cluster around zero, with a long tail of positive and negative effects. The estimated absolute average effect of a mutation is just over 2%, and the inferred DFE is shown in Figure 4.

**Table 3.** Models, and parameter estimates. The indices [0] and [1] indicate negative and positive mutation effects, respectively. 95% credible intervals are shown in square brackets.

| Model | Mean mutation effect, $\beta/\alpha$ | | Shape parameter, $\beta$ | | $q$ |
| --- | --- | --- | --- | --- | --- |
|  | [0] | [1] | [0] | [1] |  |
| Two-sided<br>gamma | 0.022<br>[0.019, 0.027] |  | 0.53<br>[0.40, 0.69] |  | 0.50<br>[0.43, 0.58] |
| Two-sided<br>gamma,<br>different<br>means | 0.023<br>[0.017, 0.030] | 0.041<br>[0.029, 0.060] | 0.53<br>[0.40, 0.70] |  | 0.51<br>[0.42, 0.58] |
| Two-sided<br>gamma,<br>different<br>means and<br>shapes | 0.023<br>[0.015, 0.032] | 0.023<br>[0.015, 0.033] | 0.53<br>[0.29, 1.0] | 0.69<br>[0.31, 1.2] | 0.49<br>[0.35, 0.64] |

**Figure 4.**
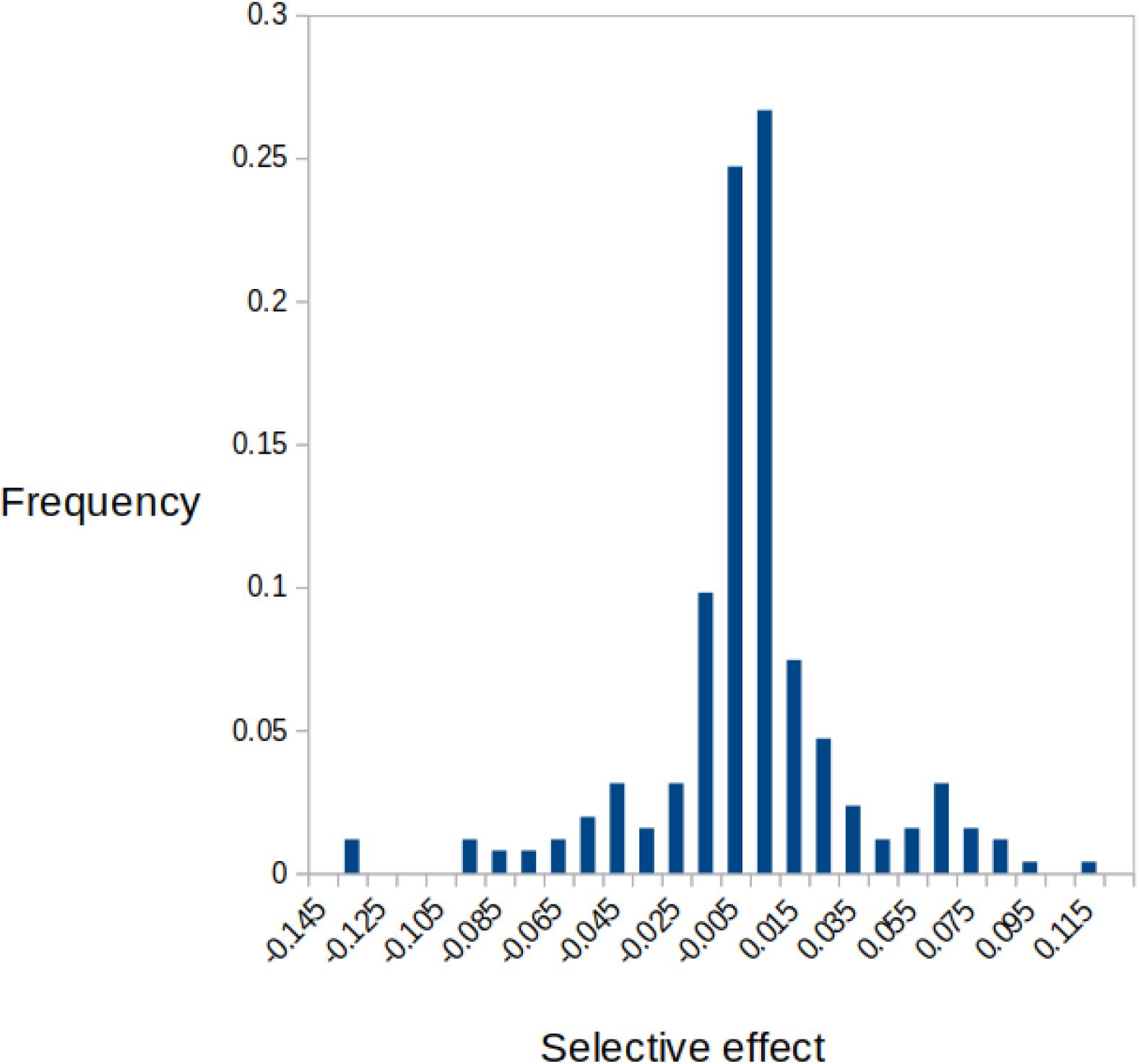
Inferred DFE, assuming the two-sided gamma distribution of effects with the same shape and scale parameters for negative- and positive-effect mutations.

### Tests for differences between annotated mutations

Using the genome annotation for *C. reinhardtii*, we tested whether certain annotated types of mutations relating to protein-coding gene function have smaller or larger effects than others by calculating the difference in mean effect or mean squared effect between them (Table S2). All differences are nonsignificant.

As a complementary analysis, we estimated the effect of mutations on annotated coding sequences using SnpEff (Cingolani et al. 2012). Of the ∼42% of mutations that could be classified using this approach, 35% were estimated to have low impact and 61% moderate impact. These classifications largely coincided with the synonymous and nonsynonymous classifications used above. Only four high impact mutations were predicted, all of which were nonsense mutations.

## Discussion

We have inferred the DFE for a set of spontaneous mutations from a MA experiment in *C. reinhardtii* by tracking their frequencies under natural selection in the laboratory and equating the observed frequency changes to the corresponding selection coefficients. The analysis of the data is made complicated by the fact that groups of mutations are linked on the same chromosome, implying that every mutation on a chromosome is expected to change in frequency even if only a subset of mutations is subject to selection. We have therefore crossed MA lines with an unmutated ancestor to obtain recombinants carrying the mutations in all possible combinations and measured frequency changes in the resulting recombinant populations after experimental evolution.

We made use of the genetic map of *C. reinhardtii* for predicting changes of frequency of linked mutations on the same chromosome. In our data, frequency is estimated on the basis of numbers of sequencing reads, and we have used this information to compute the likelihood for a set of predicted frequencies. The likelihood is then used in a Bayesian setting to compute the posterior distribution of parameters of the DFE. To our knowledge, with the exception of our previous study (Böndel et al 2019), there have been no other attempts to fit a parameterized distribution to infer the DFE for new mutations. For example, Flynn et al (2020) produced a graphical representation of the DFE based on estimates of individual mutation effects, but did not estimate the DFE’s parameters within a statistical model. This is desirable, because a simple plot of the individual effects of mutations will be inflated by sampling variance, as will summary statistics derived from this distribution.

We observed substantial changes in the frequencies of the majority of mutations, and these changes were highly repeatable among replicates starting from the same base population. This is consistent with the action of natural selection under experimental evolution. The inferred DFE is broadly consistent with that inferred in our previous study on a different *C. reinhardtii* strain (Böndel et al 2019) in which we measured growth rate and assayed genotypes of many recombinants that emerged from a cross. If we assume a two-sided gamma DFE with equivalent scale and shape parameters for positive- and negative-effect mutations, the estimated proportions of positive-effect mutations are almost identical between the two studies. The estimated shape parameter in the current study is somewhat higher than the previous study (β = 0.53 versus 0.32), implying a somewhat less leptokurtic distribution, and the mean mutational effect is about four-fold higher (0.022 *versus* 0.0049). The biological significance of these differences is unknown.

Our estimated DFE is based on certain assumptions, and there are several potential causes of inaccuracy and/or bias. First, the mutations whose frequencies we tracked were those detected by Illumina sequencing in a previous study (Ness et al 2015), but there are some mutations, including transposable element movements and large scale rearrangements, that we do not currently know about. Selection acting on these unknown mutations will therefore induce frequency changes at linked sites and generally inflate estimates of the strength of selection. Second, the linkage map for the strain we are working with may differ from the one that was assumed. Presumably any differences will lead to over/underestimation of effects, but not in a systematic way. Third, our analysis incorporates changes of frequency that occurred up to time 0 and changes that subsequently occurred up to time *t*. We do not know the number of generations up to time 0, but it is clear that in several cases mutation frequencies had already changed by then. Furthermore we are assuming a certain value for the number of generations of experimental evolution, but do not have a precise measure of this.

We infer that there is a high frequency of mutations with positive effects on fitness, which we also observed in our previous study in a different *C. reinhardtii* strain (Böndel et al 2019). Such a high frequency is a surprising finding, since it has long been argued that the majority of new mutations are likely to be neutral or deleterious (summarized by Keightley and Lynch 2003). Broadly, the majority of nucleotide sites in compact genomes such as in *C. reinhardtii* are at sites that are selectively constrained, and relatively few sites can evolve free from the influence of natural selection. In nature, the dominant force of natural selection appears to be purifying selection, since fitness in natural populations is likely to be close to an adaptive peak and most changes are therefore harmful. Direct evidence for this comes from attempts to infer the fraction of advantageous amino acid mutations, based on analysis of the site frequency spectrum (e.g., Keightley et al 2016; Tataru et al 2017). Most studies utilizing mutation accumulation also suggest that the net directional effect of mutations is negative (summarized by Halligan and Keightley 2009).

One possible explanation for the higher than expected frequency of advantageous mutations is that there was selection against deleterious mutations and in favour of advantageous mutations during the generation of the MA lines. This could have affected the strain that was the subject of the current experiment and the strain studied by Böndel et al (2019). In the experiment described here, there were approximately 11 generations of growth between each transfer in the MA experiment, where selection could operate to change the frequencies of *de novo* mutations. To investigate the influence of selection on the estimated DFE, we implemented the method recently developed by Wahl and Agashe (2021) to predict the extent of under- or over-contribution to the DFE for mutations with given selection coefficients, assuming that there is a doubling of cell number for *t* = 11 generations during mutation accumulation. Based on equation (2) of Wahl and Agashe (2021), and assuming a two-sided gamma distribution with parameters specified in Table 3, the corrected estimate for the frequency of positive effect mutations is *q* = 0.46 (uncorrected = 0.50). This suggests that selection during mutation accumulation had only a modest impact. Presumably, this is because our inferred DFE is leptokurtic, and most of the density is concentrated near zero. Furthermore, the mean absolute selective effect is relatively low (0.022), and the 11 generations of growth between transfers in the MA experiment were insufficient to lead to substantial frequency changes over the bulk of the distribution. The contrast between corrected and uncorrected DFEs (Figure 5) shows only a slight downward shift in the corrected distribution.

**Figure 5.**
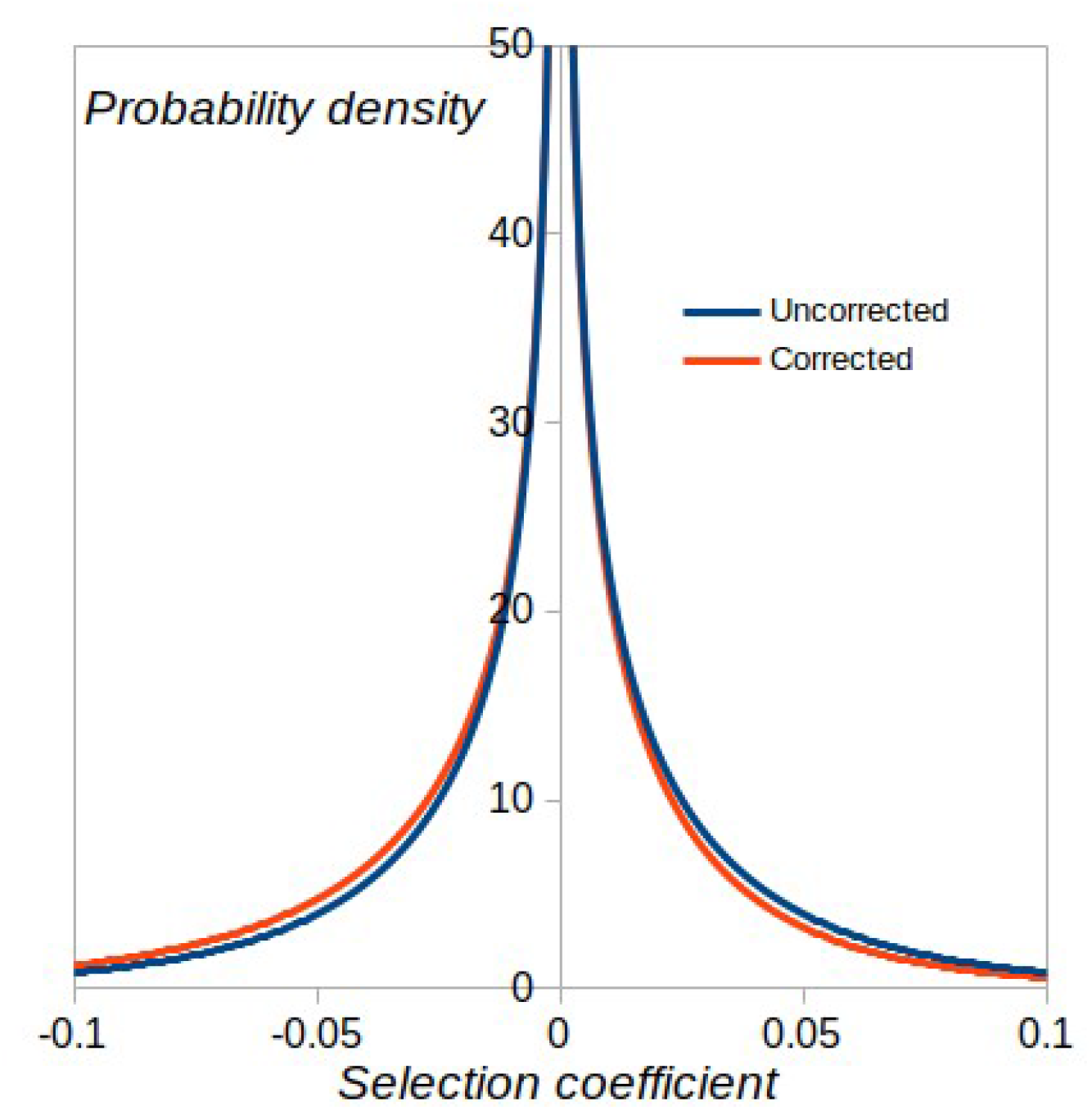
Uncorrected DFE assuming the parameters in Table 3 and corrected DFE generated by applying the method of Wahl and Agashe (2021).

Although our experimental design attempted to mitigate it, there will also have been selection against those colonies that were too small to see at the time of transfer during the MA experiment. This would most likely select against strongly deleterious mutations, but the effect of this between-colony selection is difficult to quantify.

If selection during mutation accumulation was not the explanation for the high frequency of positively selected mutations, then we must seek other explanations. Although the strains we studied were isolated three decades ago, *C. reinhardtii* can be maintained for long periods without cell division and it is probable that they have passed through insufficient generations to adapt to the novel laboratory environment. Fitness is therefore likely to be far from an adaptive peak, and therefore the fraction of advantageous mutations is expected to be higher than in a natural environment (Orr 1998). Little is known about the ecology of *C. reinhardtii*, but the laboratory clearly represents an extremely artificial environment. *C. reinhardtii* has been isolated from soil and may also be present in freshwater, where the species experiences day/night cycles, fluctuating temperatures and resource availability, biotic interactions (predators, pathogens etc.), and so on. Adaptations to life in the field include a complex metabolism (autotrophy and heterotrophy) and motility in response to light and nutrients (Sasso et al. 2018). It is plausible that loss of function mutations in genes no longer required in the laboratory are advantageous. Loss of function mutations have frequently arisen under laboratory conditions, for example certain strains have lost the ability to utilise nitrate after culture with an alternative nitrogen source (Harris 2009; Gallaher et al. 2015), although it is unknown if the underlying mutations in such cases were positively selected.

## Acknowledgements

This project has received funding from the European Research Council under the European Union’s Horizon 2020 Research And Innovation Programme (Grant Agreement no. 694212).

## Supplementary materials

### Supplementary Text A

#### Calculation of scaled recombination rates per chromosome

Let *y*_*i*_ be the scaled recombination rate for chromosome *i, x*_*i*_ be its length, and *b* the slope of the linear relationship between recombination rate and map length, i.e.,

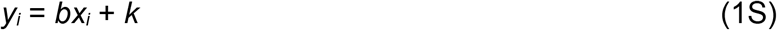

If *n* is the number of chromosomes, the mean recombination rate is:

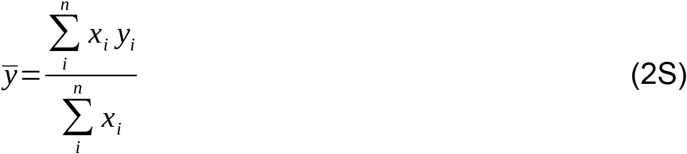

Substituting (1S) into (2S), we obtain:

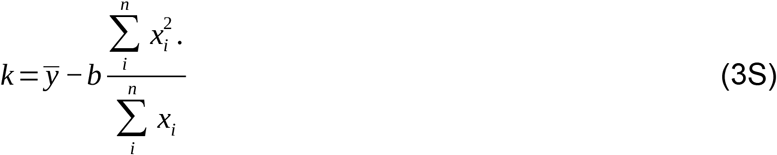

**Table S1.** Estimated numbers of generations of experimental evolution.

| Recombina<br>nt<br>population | Number of<br>days of<br>experiment<br>al evolution | Number of<br>generations<br>replicate 1 | Number of<br>generations<br>replicate 2 | Number of<br>generations<br>replicate 3 | Mean<br>number of<br>generations | SD of<br>number of<br>generations |
| --- | --- | --- | --- | --- | --- | --- |
| L06 | 42 | 58.33 | 57.92 | 57.96 | 58.07 | 0.2257 |
| L09 | 42 | 58.79 | 58.90 | 59.15 | 58.96 | 0.1879 |
| L10 | 46 | 59.95 | 59.38 | 60.80 | 60.05 | 0.7156 |
| L12 | 46 | 61.03 | 60.60 | 59.70 | 60.44 | 0.6766 |
| L13 | 46 | 59.40 | 60.02 | 60.94 | 60.12 | 0.7739 |
| L14 | 46 | 60.61 | 60.81 | 60.22 | 60.55 | 0.2968 |
| L15 | 46 | 60.10 | 61.32 | 60.61 | 60.67 | 0.6121 |

**Table S2.** Difference in effect sizes and effect sizes squared between annotation categories along with P-values obtained from 10,000 bootstraps.

| **Annotation type contrast** | **Mean effect difference (x1000)** | **P-value** | **Mean squared effect difference (x1000)** | **P-value** |
| --- | --- | --- | --- | --- |
| Exonic v non-exonic | -0.00316 | 0.91 | 0.000038 | 0.77 |
| Genic v non-genic | 0.00851 | 0.89 | 0.000009 | 0.78 |
| Nonsynonymous v synonymous | -0.0110 | 0.77 | 0.000594 | 0.71 |

| Annotation type<br>contrast | Mean effect<br>difference<br>(x1000) | P-value | Mean squared<br>effect difference<br>(x1000) | P-value |
| --- | --- | --- | --- | --- |

**Figure S1:**
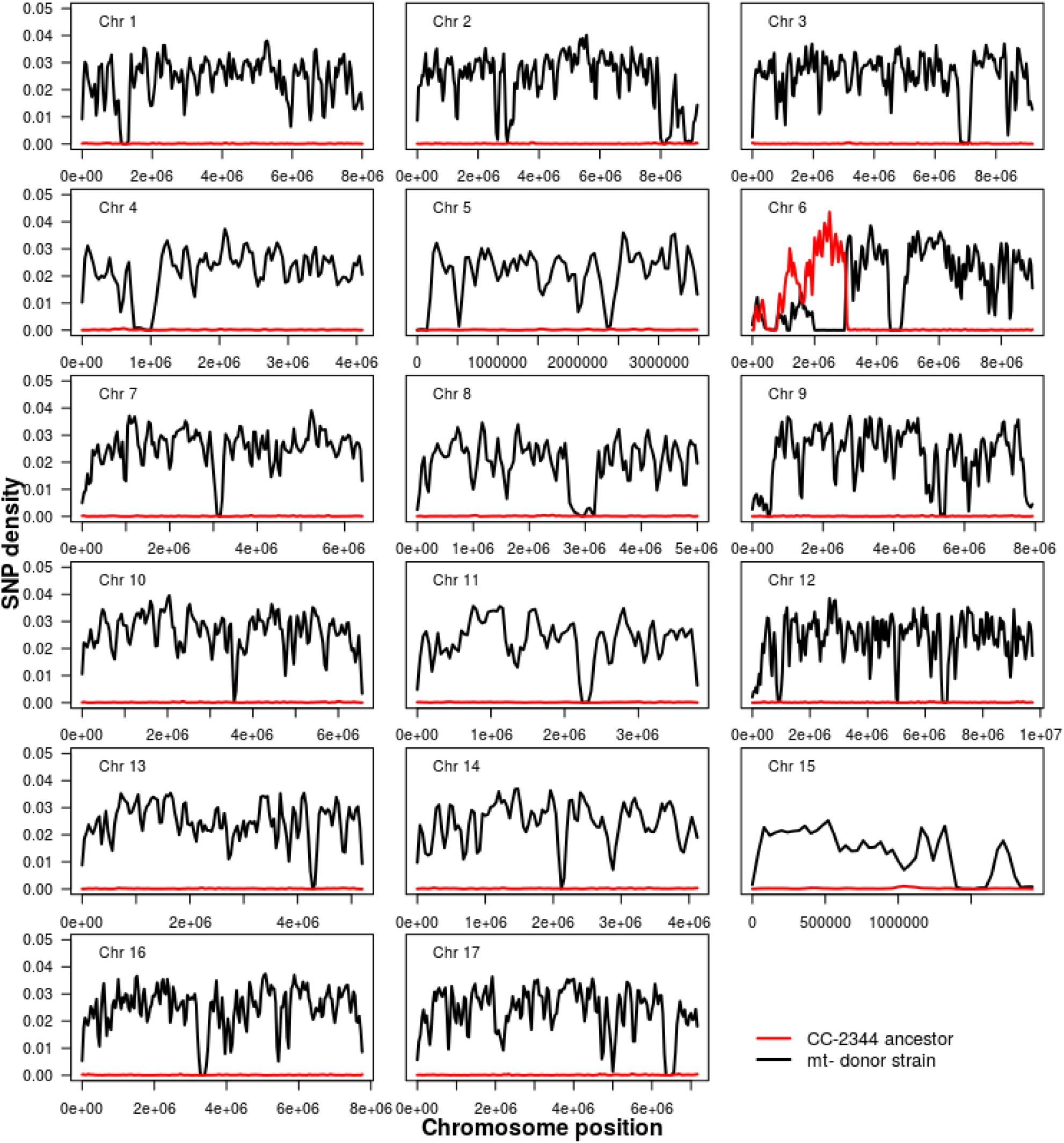
SNP densities along the 17 chromosomes between the compatible ancestor for CC-2344 and its two ancestral strains. SNP densities were calculated for 80-kb windows along the chromosomes between the compatible ancestor and CC-2344 (red) and between the compatible ancestor and the mating type *minus* (mt-) donor strain (black). A SNP density of 0 indicates no genetic differences between the compatible ancestor and the strain it was compared to.

**Figure S2.**
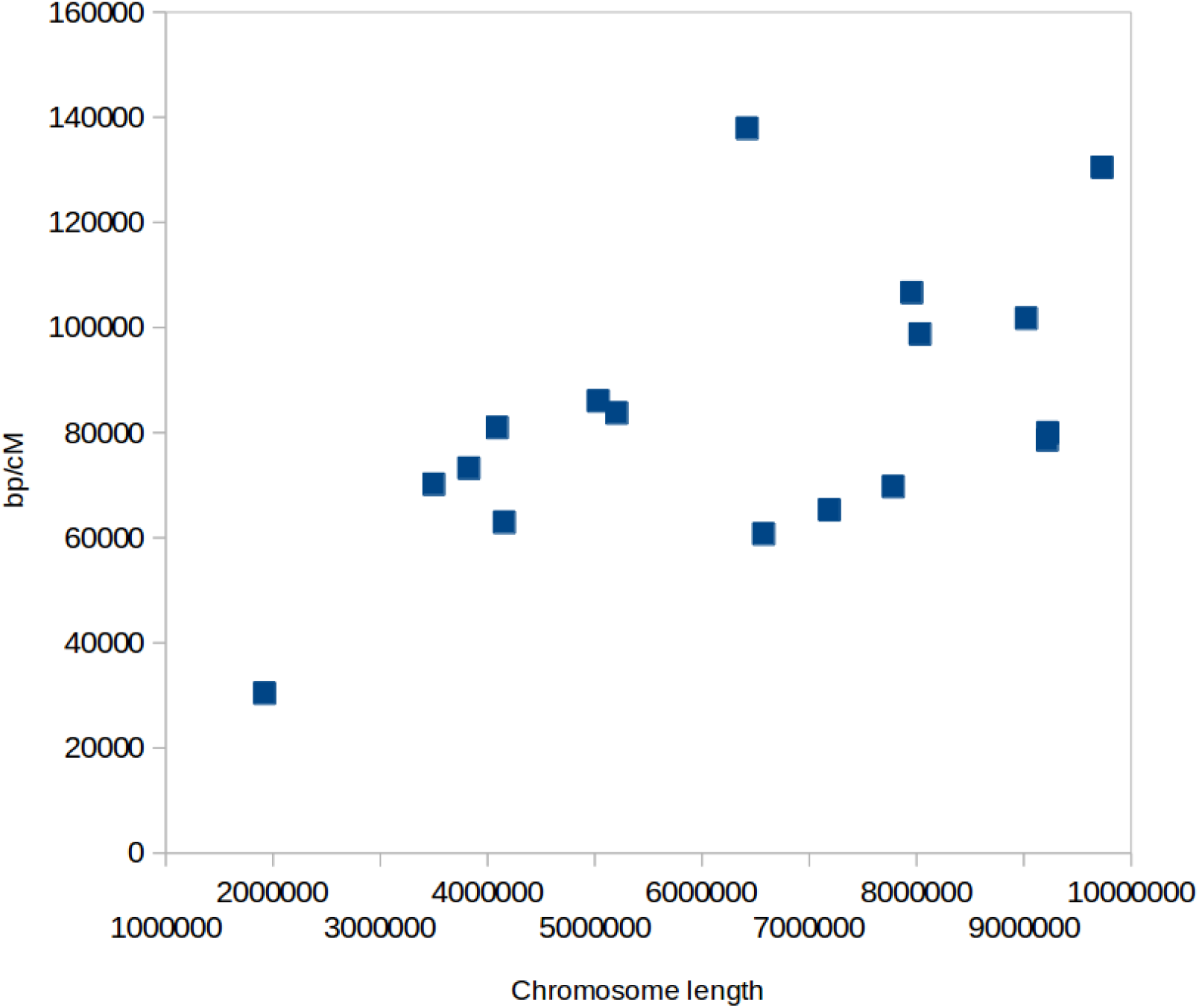
Relationship between chromosome length (in bases) and base pairs per cM, estimated from pairwise linkage disequilibrium in *C. reinhardtii* from a natural population (data from Hasan and Ness 2020). (Pearson correlation *r* = 0.56; linear regression: bp/cM = 0.00616 x length + 43,900, *P* = 0.015).

**Figure S3.**
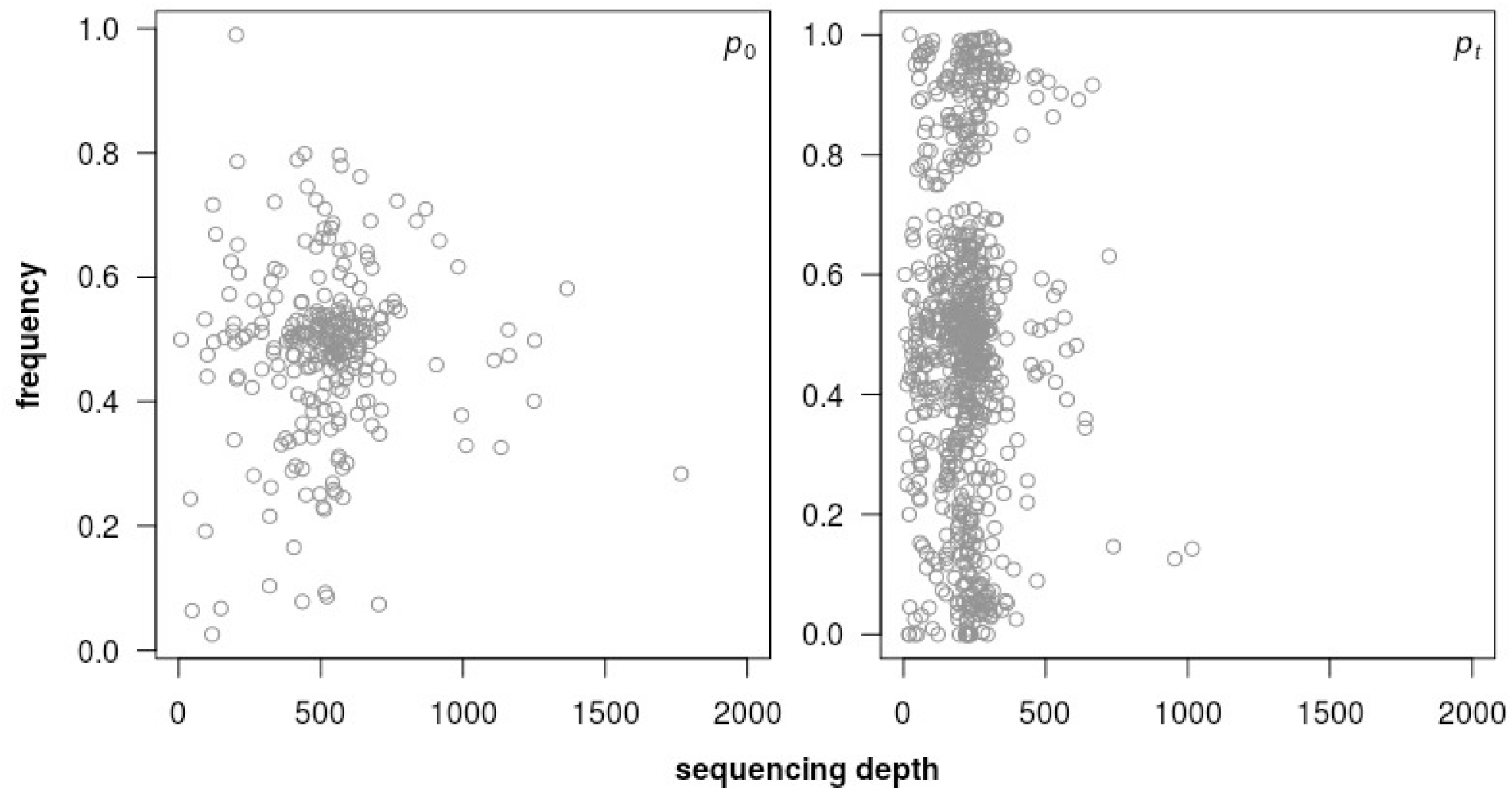
Mutation frequencies *versus* sequencing depth at times 0 and *t*. Mean sequencing depth for the mutations are 520.7x and 221.6x for times 0 and *t*, respectively.

**Figure S4.**
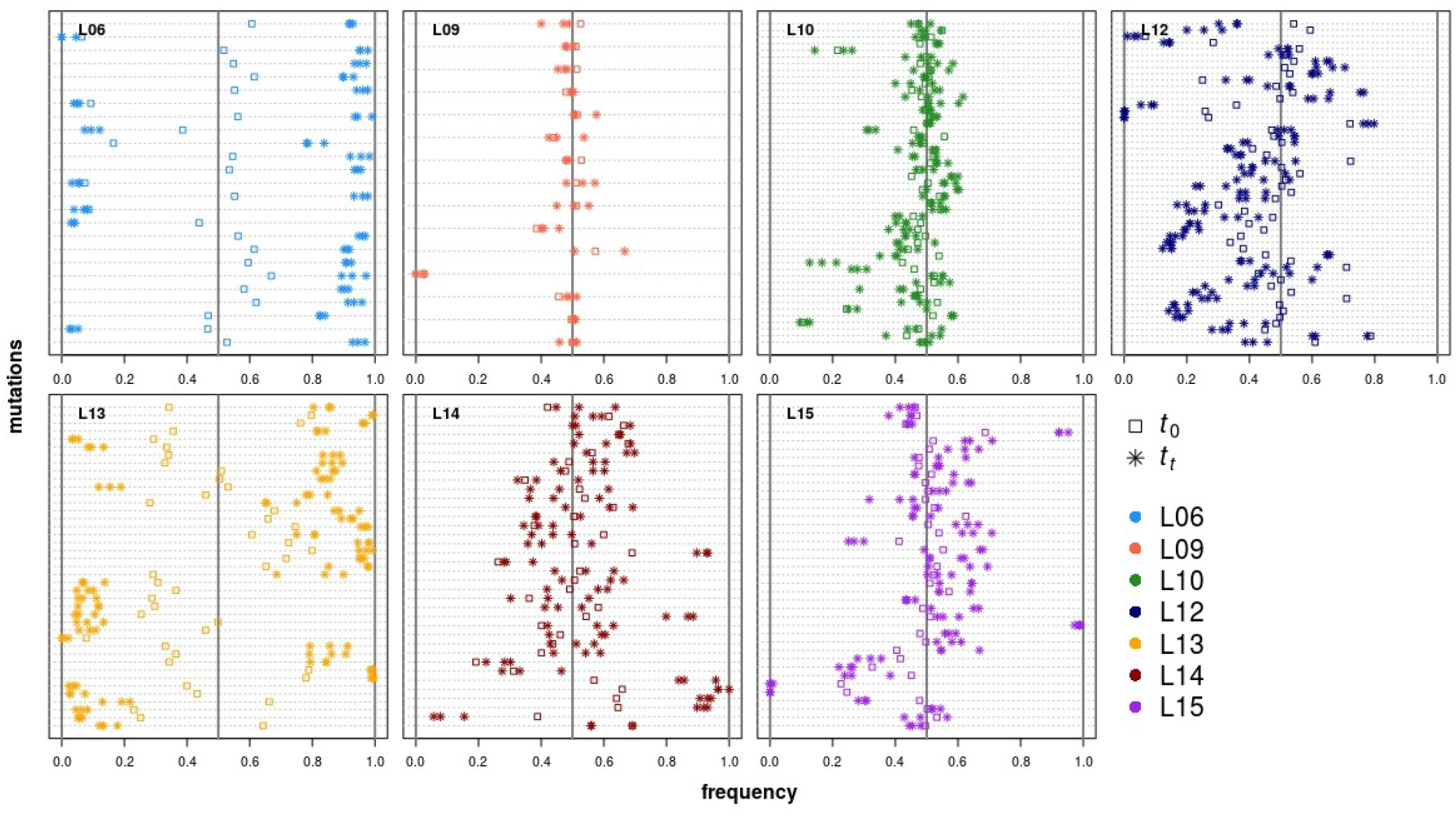
Frequencies of the individual mutations before and after experimental evolution. Mutations of each MA line are shown from top to bottom in the order in which they occur in the genome. Squares denote the mutation frequencies at *t*_0_, and stars denote the mutation frequencies of the three replicates at *t*_*t*_. The different MA line recombinant pools are shown in the different panels.

**Figure S5:**
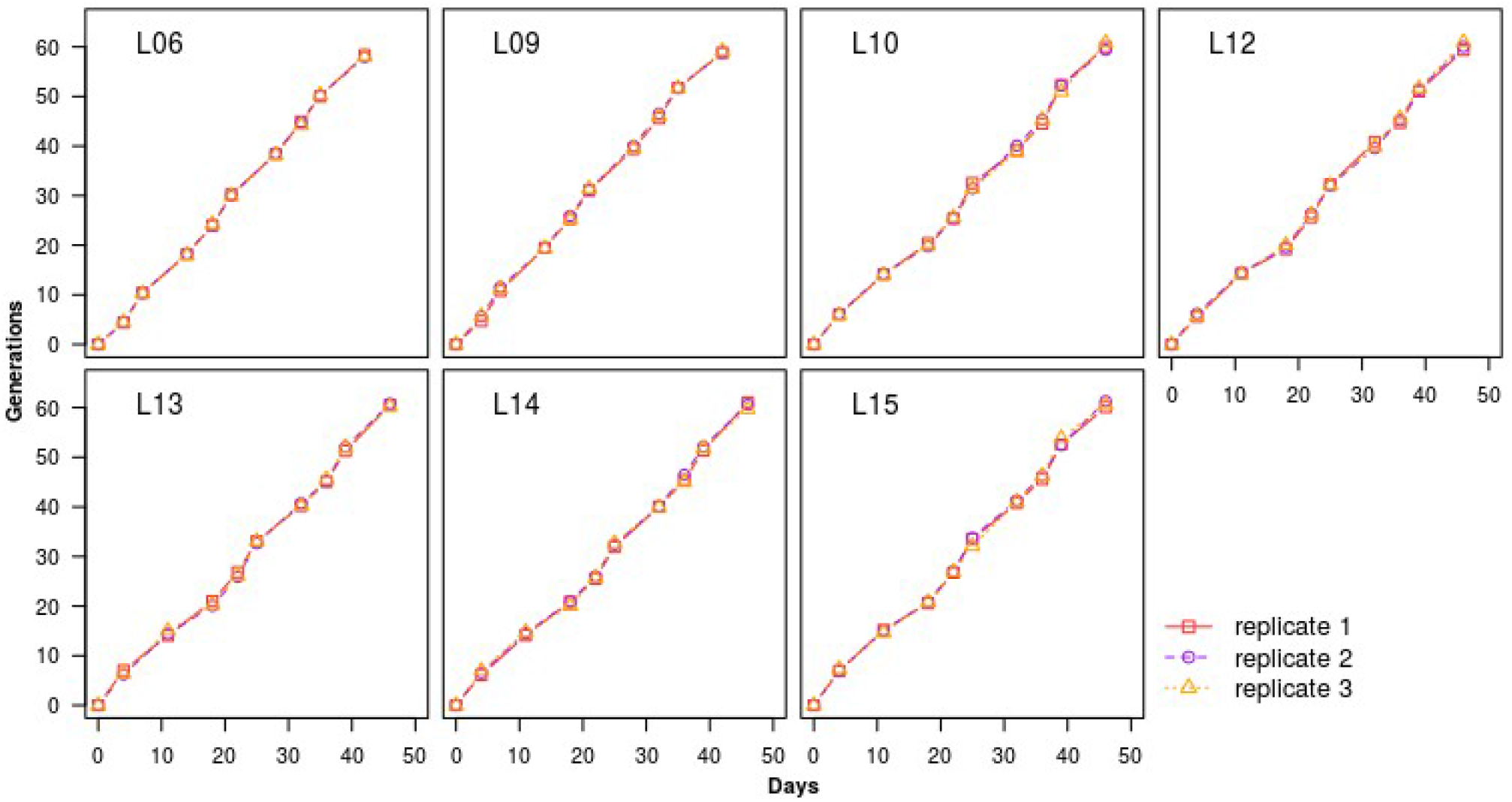
Increase of generations during the course of the experiment. Each panel shows the cumulative number of generations of the three replicates of each of the seven recombinant pools derived from a backcross between MA line and compatible ancestor. Number of generations was estimated from OD measurements conducted at each transfer.

**Figure S6.**
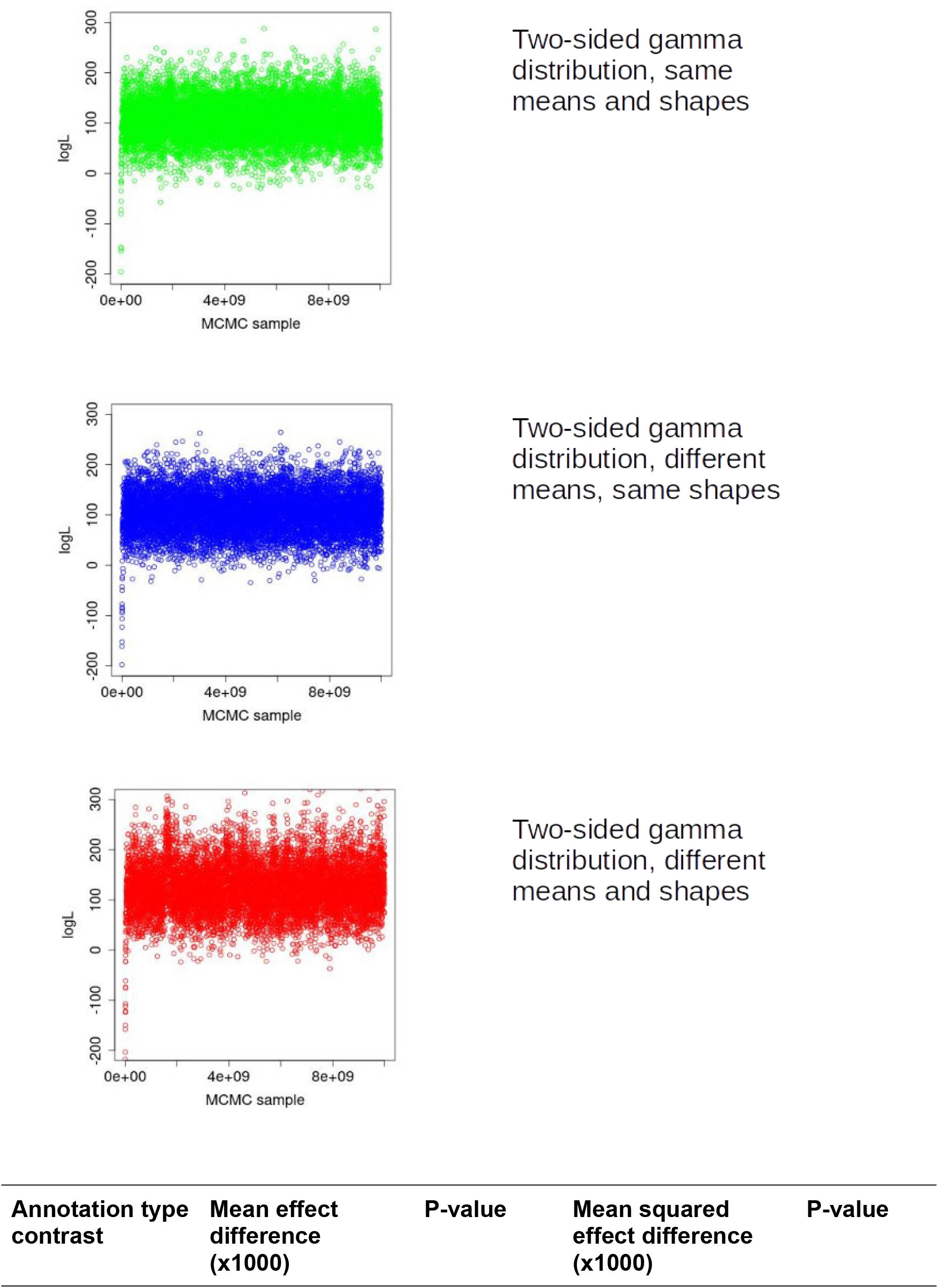
Output of MCMC sampler for three DFEs.

## References

Armstrong, J., Hickey, G., Diekhans, M., Fiddes, I. T., Novak, A. M., Deran, A., … Paten, B. (2020). Progressive Cactus is a multiple-genome aligner for the thousand-genome era. Nature 587: 246–251.

Bold HC (1942). The cultivation of algae. The Botanical Review 8: 69–138.

Böndel, K. B., Kraemer, S. A., Samuels, T. S., McClean, D., Lachapelle, J., Ness, R. W., Colegrave, N. and Keightley, P. D. (2019). Inferring the distribution of fitness effects of spontaneous mutations in Chlamydomonas reinhardtii. PLoS Biology 17: e3000192.

Chen, J, Glémin, S & Lascoux, M. (2017). Genetic Diversity and the Efficacy of Purifying Selection across Plant and Animal Species. Molecular Biology and Evolution doi:10.1093/molbev/msx088.

Cingolani, P., Platts, A., Wang, L. L., Coon, M., Nguyen, T., Wang, L., Land, S. J., Lu, X. and Ruden, D. M. (2012). A program for annotating and predicting the effects of single nucleotide polymorphisms, SnpEff: SNPs in the genome of Drosophila melanogaster strain w1118; iso-2; iso-3. Fly (Austin) 6: 80–92.

Craig, R. J., Hasan, A. R., Ness, R. W., & Keightley, P. D. (2021). Comparative genomics of Chlamydomonas. Plant Cell 33: 1016–1041.

DePristo MA, Banks E, Poplin R, Garimella KV, Maguire JR, Hartl C, Philippakis AA, del Angel G, Rivas MA, Hanna M, et al. (2011). A framework for variation discovery and genotyping using next-generation DNA sequencing data. Nat Genet 43: 491–498.

Eyre-Walker, A. and Keightley, P. D. (2007). The distribution of fitness effects of new mutations. Nature Reviews Genetics 8: 610–618.

Eyre-Walker, A. and Keightley, P. D. (2009). Estimating the rate of adaptive molecular evolution in the presence of slightly deleterious mutations and population size change. Molecular Biology and Evolution 26: 2097–2108.

Flynn, Julia, Rossouw, Ammeret, Cote-Hammarlof, Pamela, Fragata Inês, Mavor, David, Hollins, Carl, Bank, Claudia Bolon, Daniel. (2020). Comprehensive fitness maps of Hsp90 show widespread environmental dependence. eLife 9:e53810.

Gallaher SD, Fitz-Gibbon ST, Glaesener AG, Pellegrini M, Merchant SS. 2015. Chlamydomonas genome resource for laboratory strains reveals a mosaic of sequence variation, identifies true strain histories, and enables strain-specific studies. Plant Cell 27: 2335–2352.

Halligan, D. L. and Keightley, P. D. (2009). Spontaneous mutation accumulation studies in evolutionary genetics. Annual Review of Ecology, Evolution and Systematics 40: 151–172.

Harris EH. 2009. The Chlamydomonas Sourcebook (Second Edition): Introduction to Chlamydomonas and Its laboratory use. Academic Press.

Hasan, A. R. and Ness, R. W. (2020). Recombination rate variation and infrequent sex influence genetic diversity in Chlamydomonas reinhardtii. Genome Biology and Evolution 12: 370–380.

Hickey, G., Paten, B., Earl, D., Zerbino, D., & Haussler, D. (2013). HAL: a hierarchical format for storing and analyzing multiple genome alignments. Bioinformatics, 29: 1341–1342.

Johnson, M. S.,Martsul, A., Kryazhimskiy, S. and Desai, M. M.. 2019. Higher-fitness yeast genotypes are less robust to deleterious mutations. Science 366: 490–493.

Keightley, P. D. and Lynch, M. (2003). Towards a realistic model of mutations affecting fitness. Evolution 57: 683–685.

Keightley, P. D., Campos, J. L., Booker T.R. and Charlesworth, B. (2016). Inferring the frequency spectrum of derived variants to quantify adaptive molecular evolution in protein-coding genes of Drosophila melanogaster. Genetics 203: 975–984.

Li H, Durbin R. (2009). Fast and accurate short read alignment with Burrows–Wheeler transform. Bioinformatics 25: 1754–1760.

Li H, Handsaker B, Wysoker A, Fennell T, Ruan J, Homer N, Marth G, Abecasis G, Durbin R. (2009). The Sequence Alignment/Map format and SAMtools. Bioinformatics 25: 2078–2079.

Liu, H. et al. (2018). Tetrad analysis in plants and fungi finds large differences in gene conversion rates but no GC bias. Nat Ecol Evol. 2:164–173.

McDonald, MJ, Daniel P. Rice 1,2 * & Michael M. Desai. 2016. Sex speeds adaptation by altering the dynamics of molecular evolution. Nature 531: 233–236.

McKenna A, Hanna M, Banks E, Sivachenko A, Cibulskis K, Kernytsky A, Garimella K, Altshuler D, Gabriel S, Daly M, et al. 2010. The Genome Analysis Toolkit: a MapReduce framework for analyzing next-generation DNA sequencing data. Genome Res 20: 1297– 1303.

Merchant SS, Prochnik SE, Vallon O, Harris EH, Karpowicz SJ, Witman GB, Terry A, Salamov A, Fritz-Laylin LK, Maréchal-Drouard L, et al. (2007). The Chlamydomonas genome reveals the evolution of key animal and plant functions. Science 318: 245–250.

Morgan AD, Ness RW, Keightley PD, Colegrave N. (2014). Spontaneous mutation accumulation in multiple strains of the green alga, Chlamydomonas reinhardtii. Evolution 68: 2589–2602.

Ness RW, Morgan AD, Colegrave N, Keightley PD. (2012). Estimate of the Spontaneous Mutation Rate in Chlamydomonas reinhardtii. Genetics 192: 1447–1454.

Ness RW, Morgan AD, Vasanthakrishnan RB, Colegrave N, Keightley PD. (2015). Extensive de novo mutation rate variation between individuals and across the genome of Chlamydomonas reinhardtii. Genome Research 25: 1739–1749.

O’Donnell, S., Chaux, F., & Fischer, G. (2020). Highly contiguous Nanopore genome assembly of Chlamydomonas reinhardtii CC-1690. Microbiol Resour Announc, 9: e00726–00720.

Ohta, T (1973). Slightly deleterious mutant substitutions in evolution. Nature 246: 96–98.

Orr, H. A. (1998). The Population Genetics of Adaptation: The distribution of factors fixed during adaptive evolution. Evolution 52: 935–949.

Raftery AE. (1995). Bayesian model selection in social research. Sociological Methodology 25: 111–163.

Sager R, Granick S. (1954). Nutritional control of sexuality in Chlamydomonas reinhardtii. The Journal of General Physiology 37: 729–742.

Salomé, P. A., & Merchant, S. S. (2019). A Series of fortunate events: Introducing Chlamydomonas as a reference organism. Plant Cell 31: 1682–1707.

Sasso S, Stibor H, Mittag M, Grossman AR. 2018. From molecular manipulation of domesticated Chlamydomonas reinhardtii to survival in nature. eLife 7.

Tataru, P., Mollion, M., Glémin, S. and Bataillon, T. (2017). Inference of distribution of fitness effects and proportion of adaptive substitutions from polymorphism data. Genetics 207: 1103–1119.

Wahl, L. M. and Agashe, D. (2021). Selection bias in mutation accumulation. Biorxv.

